# MoonFit, a minimal interface for fitting ODE dynamical models, bridging simulation by experimentalists and customization by C++ programmers

**DOI:** 10.1101/281188

**Authors:** Philippe A. Robert, Henrik Jönsson, Michael Meyer-Hermann

## Abstract

The modelling of biological systems often consists into differential equation models that need to be fitted to experimental data. During this complex process, the practical experience of the biologist and the theoretical abstraction of the modeller require back-and-forth refinements of the model, design of new experiments and inclusion of more data-points into the fitting procedure. Available optimization interfaces rarely simultaneously allow customizations by the programmer and the capacity for the biologist to perform simulations or optimizations with a simple interface.

Here, we provide the C++ code of a graphical user interface based on a user defined minimal C++ ODE model class. The graphical interface allows to perform simulations and optimizations without any knowledge in programming. The code was designed minimal and modular to be easily modified, with maximal freedom to link customized optimization libraries, solver or hand-made scripts. Moonfit is powerful enough to fit and compare models with high dimensionality, multiple datasets, to automatize optimizations, and to perform iterative fittings using data interpolation. We believe this will ease the interaction between modellers and experimental partners.

**Availability:** Moonfit is freely available via c++ source code and accompanying scripts from gitlab.com/Moonfit/MoonLight.

## 1 Introduction

The dynamics of biological systems can often be represented as ordinary differential equations (ODEs), meaning equations describing the time-evolution of observed variables such as populations of cells, transcription factors, organ size, signaling strength [1]. Biological mechanisms are translated into ODEs describing how each variable impact each-other, such as activations or inhibitions, and the strength and complexity of these mechanisms are usually unknown parameters. Only the time evolution of the variables is usually measured in different conditions, and one would want to extract the unknown parameters values from experimental datasets. This is called ‘parameter estimation’ or ‘fitting’, and consists into performing a huge number of simulations with different parameter values, comparing each simulation to the data, and keeping the parameter set with the lowest difference (cost) to the data [2]. Depending on the quality of the best simulation, the model is either validated (success) or requires to be modified (reject) in order to accurately describe the data. A validated model can be used to make predictions while the iterative redesign of non-fitting models gives understanding on the biological mechanisms required to explain the complexity of the data. Therefore, parameter estimation is of high relevance for biological modelling [3, 4].

With the formulation of a cost function between experimental data and respective simulations, a fitting can be seen as a ‘black box’ optimization problem, with a list of *N*_*P*_ parameters and a cost function to minimize. Optimization techniques are very diverse, and their efficiency is highly dependent on the type of problem and the properties of the *N*_*P*_ dimensional landscape of the cost []. Gradient-based optimizers compute the local derivatives of the cost function along all dimensions, and are suited to perform local optimization, such as the steepest descent method, fmincon (Matlab) or Newton-derived methods [5]. These methods need to be coupled to a global strategy such as multistart or combined with a global optimizer to account for global optima. Derivative-free optimizers include Particle Swarm [6], Simulated Annealing [6], or Evolutionary Algorithms, that emulate an artificial natural selection on a population of parameter sets, such as SRES [7], CMA-ES [8] or differential evolution [9].

Parameter optimization can be an essential step for the successful development of systems biology models. Different optimizers with different options need to be compared until a good quality of fitting is reached. Popular systems biology toolboxes such as COPASI [10], MEIGO [11], PESTO [12] and D2D [13] conveniently include a large choice of optimizers under the same syntax. However, the multitude of quantitative data coming available at the molecular and cellular resolution makes it almost impossible to create a static tool that can incorporate all possible data sets for model optimization, and more dynamic and adjustable tools are required.

Further, judging the quality and biological significance of a fitting most often requires the sharp eye of an experimentalist and the capacity to interact with the model by adjusting parameter sets. Several powerful user interfaces allow to simulate and optimize ODE networks. The most famous are probably Cell Designer to formulate biochemical or cell populations networks [14] and COPASI to optimize them to experimental data [10]. CellNetAnalyzer allows in particular to optimize metabolic networks [15] and SBPOP / IQM tools in Matlab (MathWorks Inc, MA) allows to simulate and optimize ODEs automatically generated from a pseudo-code definition of the equations [16]. These powerful tools give the opportunity to experimentalists to develop and fit simple and more complex models with limited knowledge in programming.

In the context of collaborations between modellers and experimentalists, the modeller might want more flexibility regarding the model formulation or the capacity to use other tools than the built-in solvers or optimizers, and might want to develop home made scripts or cost functions that are more suited to the experimental data. From a programming point of view, it is quite straightforward to implement ODEs and use solvers or optimizers from libraries, but endorsing the experimentalist with the capacity to monitor fittings by him/herself would be much simpler with a graphical interface. Here, we present MoonFit, a tool developed in c++ with the intention to bridge the gap between a core source code for simulating and optimizing ODE models adaptable to a variety of models, solvers, optimization methods and experimental input data, and performing the resulting optimization via a user friendly graphical interface.

## 2 Results

### Running a model optimization

MoonFit is a program developed in C++ to solve and optimize ODE dynamical models from a graphical interface. Only the minimal information of the ODEs and the design of simulations have to be defined inside a C++ class following a predefined structure. Then, the provided graphical interface directly gets information from the model class and runs simulations and parameter optimizations from this model by choosing parameter boundaries and combinations of parameters to optimize. Full freedom is given on the implementation of the model class and equations as long as it follows the predefined structure. Further, we show how to replace the default solver or optimizers by another C++ library. Basic examples associated with the code include Lotka-Volterra prey-predator dynamics, 3-stage cell development (later presented in Figure 3), and viral dynamics [17], and can directly be run to teach parameter optimization to students without programming knowledge.

### Structure of the code

The structure of the framework is shown in Figure 1. Each file corresponds to a class/module, and has been designed as simple as possible in order to be easily modifiable. For instance, different solvers, scripts or optimizer can be linked to the code (see section 4).

**Figure 1:**
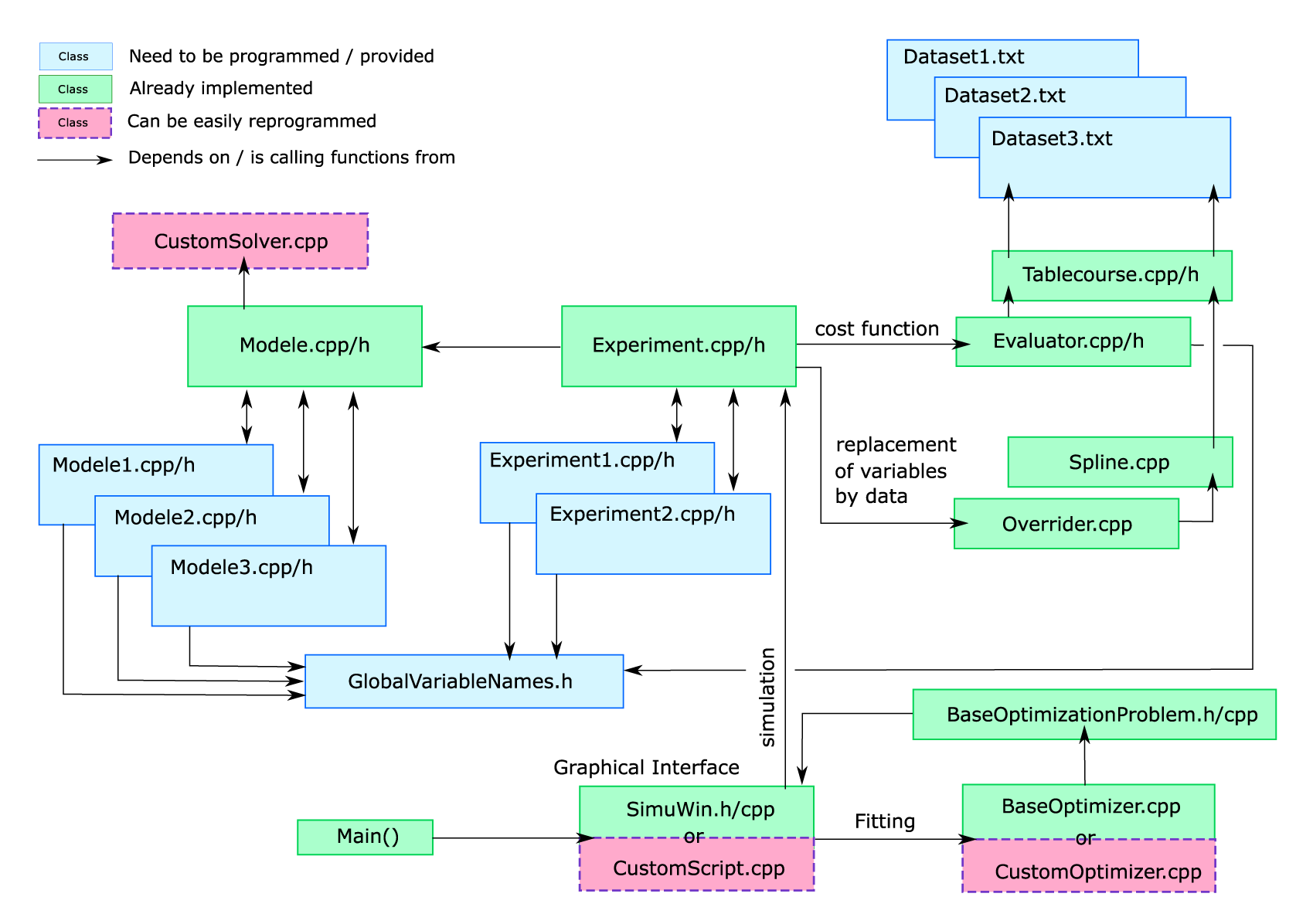
Modular structure of the code. Blue boxes: classes that need to be implemented. Dashed orange boxes: classes that can easily be replaced by another library or program at wish. Arrows A –> B: the class A can call or get informations from B. The ODEs and parameter boundaries need to be defined in sub-classes of the Modele mother class. The Modele class has built-in functions to solve the ODEs and run simulations according to a certain parameter set. The experiments to be simulated are defined as sub-classes of the mother class Experiment. The Evaluator class is used to load experimental datapoints from text-files, either manually or as tab-separated text files using the tablecourse class. Each time the Experiment class is asked to perform a simulation, it uses the chosen Model subclass to initializes and run a simulation, it stops simulation at each time-point requested by the evaluator, saves the simulated value and ultimately returns the cost of this simulation compared to the data. The Experiment class also permits to replace certain variables by their interpolated curve from data (linear or 3-splines) by using the overrider and spline classes, allowing to perform iterative fittings. The class manageSims (without GUI) or simuWin (GUI) provide functions to perform optimizations, sensitivity or identifiability by simulating the experiments and retrieving their cost. The classes manageSims and simuWin are formatted as a ‘baseOptimizationProblem’, meaning a black-box function/class that returns a cost from a N-dimensional parameter set. Therefore, it can easily be given to any optimizer library. The default optimizer provided is called BaseOptimizer, and includes Simulated Annealing, Stochastic Ranking Evolutionary Strategies, and Classical Evolutionary Programming according to 12 different cross-overs and 8 different mutations [6].

### Graphical interface for simulations / optimizations

Basic optimizations can be started via the graphical user interface (Figure 2). It combines all the ingredients necessary for a fitting in a way that any person without programming knowledge can perform parameter optimization. It includes the choice of which parameter to optimize and their boundaries, the visualization of the simulation versus data for each variable and experiment, the capacity to replace variables by their interpolated experimental data, and the choice of different parameter sets resulting from an optimization.

**Figure 2:**
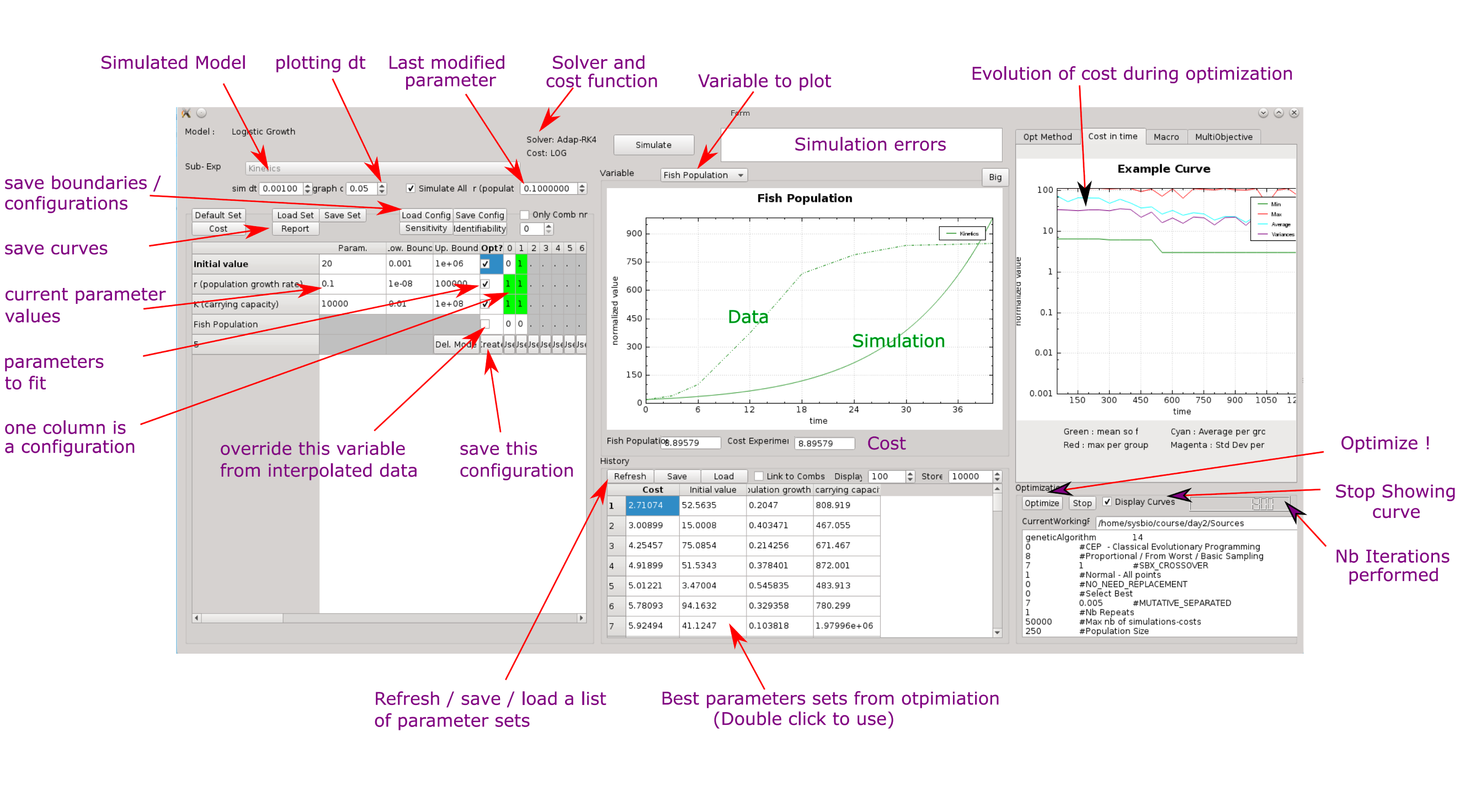
User interface. According to the defined models and experiments subclasses, the user interface allows to run simulations and perform optimizations. For a simulation, the model and a set of parameters have to be chosen. The simulation results is shown in the middle panel. For an optimization, the list of parameters to optimize, their boundaries, and which variables are simulated or replaced by interpolated data are defined as ‘configurations’. Each configuration can be optimized according to the optimizer chosen in the ‘optMethod tab’. The cost evolution over time is displayed, together with the list of best parameter sets and their cost. Clicking on a parameter set launches a simulation according to its values. Further, parameter values can be changed manually by using the last modified parameter input box.

## 3 Step-by-step definition and fitting of an ODE model

Figure 1 shows the structure of the framework, as well as the interplay between user-defined files and the provided files. Three kinds of files have to be provided for a new project: 1/ the ODEs inside model files, 2/ the experimental layout inside experiment files, and 3/ experimental data as text files. An additional file defining the name of the variables in the experimental data (GlobalVariableNames) is recommended, such that each model could simulate different variables and link their internal variables to their ‘official experimental name’.

### 3.1 A biological example

**Motivation.** Throughout the next sections, a small biological problem requiring parameter optimization will be developed. Based on the work of Bertocci et al., deficient mice for a specific gene, KLHL6, were phenotyped for the development of B cells in the spleen [19]. Three populations were considered: two transitional stages, T1 and T2 and a mature phenotype, Follicular B cell. The deficient mice show a different dynamics of development, with a strong reduction of all populations from the T1 stage. We would like to know which mechanism is/are impacted by the mutation. The authors propose that both the death of T1 cells and the conversion of T1 into T2 cells is impacted and it would be interesting to know whether the quantitative dynamics of the experimental data are consistent with this hypothesis.

The transitions between each population stage is depicted in Figure 3, together with the comparison of cell numbers between WT and deficient mice, and the differential equations to represent the biological system. The mechanisms considered are the inflow of T1 cells (*K*_*F*_), the transition between a population and the next stage (*K*_1–2_, *K*_2–3_, *K*_3–*out*_), and the death rates (*K*_*D*1_, *K*_*D*2_, *K*_*D*3_). A phenomenon of space constraint can be modelled by a negative feedback from the cells present in the compartment, inhibiting the inflow of cells. Since the major population are the mature cells, the feedback can be approximated as coming only from mature cells, and is implemented as a Michaelis-Menten dynamics, with maximal inhibition *Mi* and typical range of action *Ki*. The proposed hypothesis that KLHL6 deficiency impacts both *K*_1 − 2_ and *K*_*D*1_ can be modelled by having different values *K*_*D*1*mut*_ *K*_1–2*mut*_ in the deficient background while keeping the same values for the other parameters between both backgrounds. Note: the development of T1 cells starts around day 11 during development and the datapoint *t* = 11 with empty populations of T1, T2 and T3 can be added. In the next paragraphs, we show how to fit the experimental values to both models using MoonFit.

**Figure 3:**
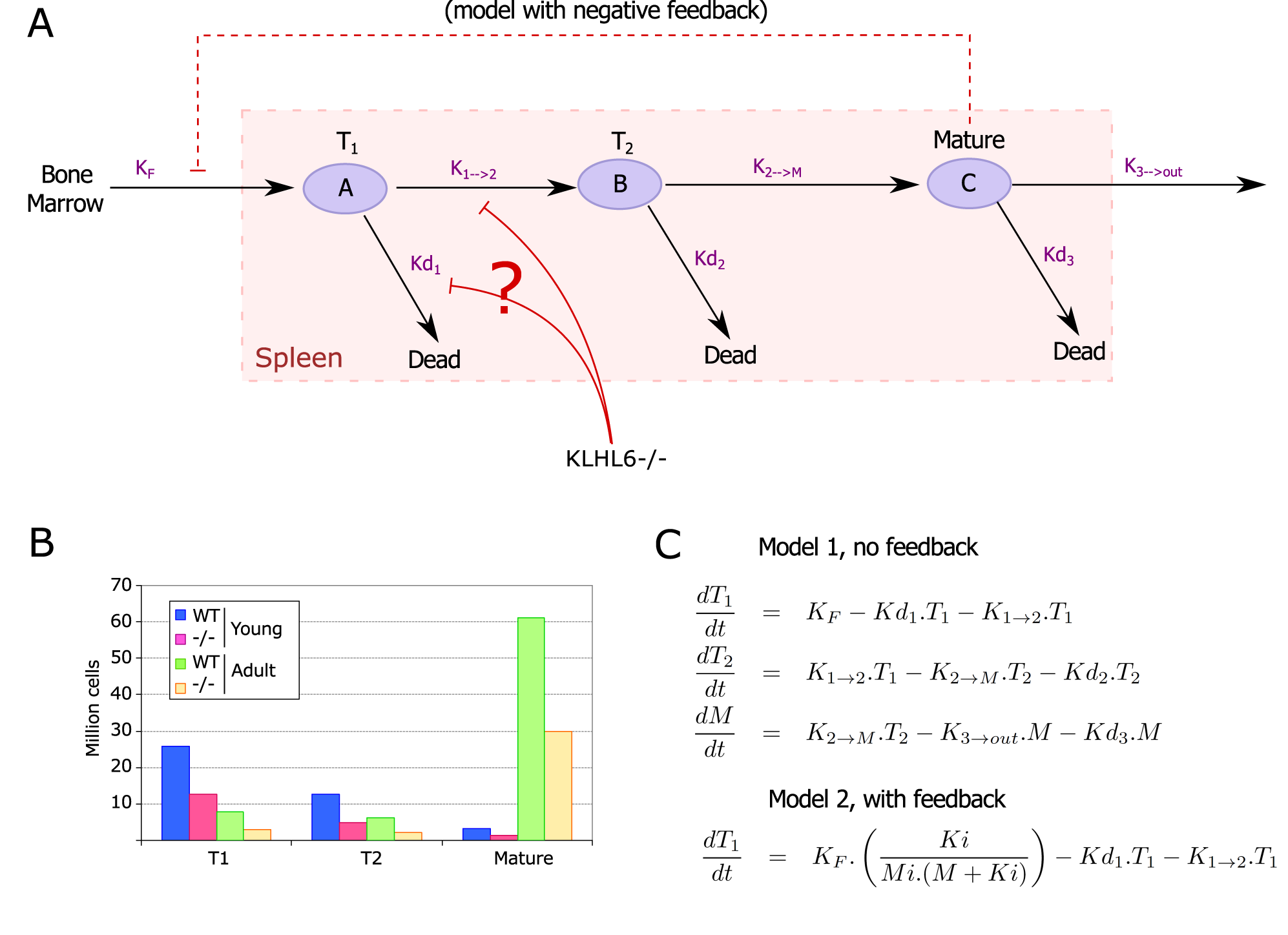
Biological example for parameter estimation. Model for the development of B cells in the spleen and experimental profile of WT and deficient mice for KLHL6. A. Developmental scheme of B cells in the spleen, with potential niche effect of mature cells inhibiting the inflow of T1 cells.B. Experimental cell numbers for the three populations at 14-20 days (young) or adult mice, as an approximation to steady-state [19] C. Associated ODE model, with or without feedback.

**Formulation of the problem as a fitting.** While parameter estimation is conceptually simple, it requires to define many technical details and can be labour intensive. After the biological question is settled, the following ingredients need to be defined:

1. The mathematical formulation of the model(s). The ODEs describing the time-evolution of the variables need to be designed, with a number *N*_*V*_ of variables and *N*_*P*_ of unknown parameters.
2. Experimental data, i.e. a list of quantitative values for some of the variables at different timepoints and under different conditions.
3. An in silico experimental layout, describing which simulations should be preformed to be compared to the experimental data: a number *N*_*C*_ of different conditions to be simulated, including the initial values for each variable, the duration of simulation and the perturbations applied to the system during the experiment.
4. Realistic boundaries for each parameter, based on biological knowledge, or very large if unknown.
5. An objective, or cost function, to be used to compare a simulation to the dataset. Here, two cost functions are provided: the rooted sum of squared differences (RMS) or the sum of log differences. If standard deviation is provided with the datasets, the cost can be normalized by the standard deviation.
6. A solver, i.e. a program to simulate the time-evolution of the variables according to the ODEs. The program proposes to use adaptive Runge-Kutta 45, or adaptive euler solvers. It has to be noted that usual solvers are designed for high precision of simulation, and become slow or can freeze when the tested parameter is far from reality. A stopping condition and a minimal dt are included when calling the solver in order to discard diverging simulations as early as possible. A cost penalty is raised for diverging simulations that increases if the divergence happens early, and the penalty is added to the data-associated cost of the parameter set.
7. An optimizer, i.e. a program or heuristic that chooses which parameter sets to simulate in order to find an optimal parameter set with minimal cost. The included optimizers are: Simulated Annealing, Stochastic Ranking Evolutionary Strategies, and Classical Evolutionary Programming according to 12 different cross-overs and 8 different mutations [6].

In the following section we explain how to set up each of these ingredients using Moonfit.

### 3.2 the ODE model

The structure of fields that are common between any different models are already programmed inside the Modele.cpp/h class, for instance the fields for saving the parameters names, or their boundaries. A new ODE model should be programmed as a sub-class of the Modele class, thereby giving it access to all fields and functions of the mother class. In that way, the definition of a new model is reduced to its minimum. All information that is specific to this model has to be written in the sub-class, namely the differential equations, the list of variables and parameters. For instance, a model could refer to one mouse strain, and different models could be built according to different mouse backgrounds or different model assumptions. As an illustration, the main fields of the mother class Model are shown in Figure 4, and a more detailed list of already implemented functions such as simulate are shown in Figure 6.

**Figure 4:**
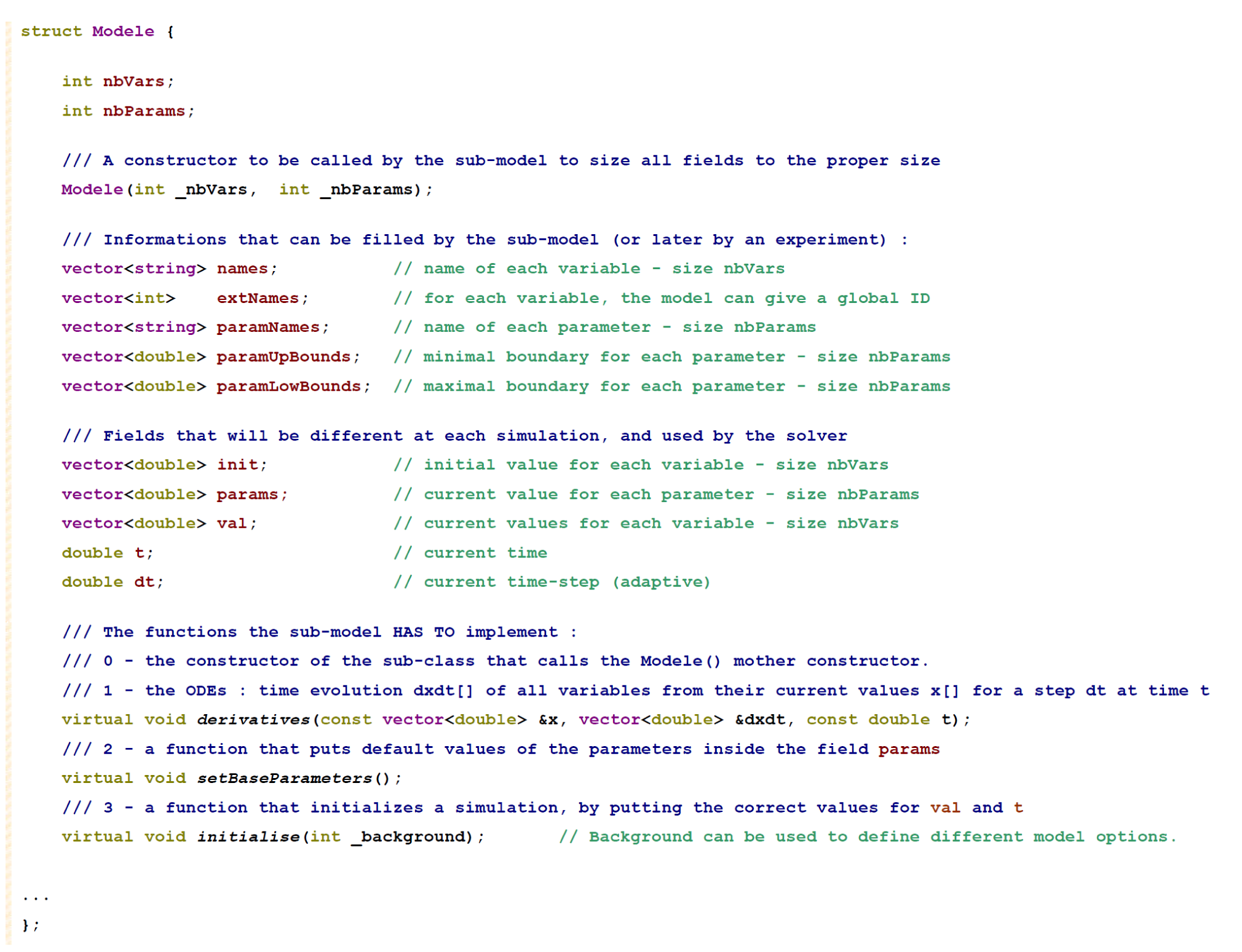
The main fields of the Modele class that will be inherited to the developed ODE models. Each ODE model will be a sub-class of the Modele mother class (’sub-model’) and can use the fields of the mother class to define the parameter names, variable names, parameter boundaries, and more. The only constraint a sub-model should fulfill is to define a constructor and three functions, namely the implementation of the derivatives (ODE) from the state vector x at time t, a function to put default parameter sets and a function to initialize a simulation according to the current set of parameters. Any sub-model following these rules will directly be usable from the graphical interface for simulations.

**Figure 5:**
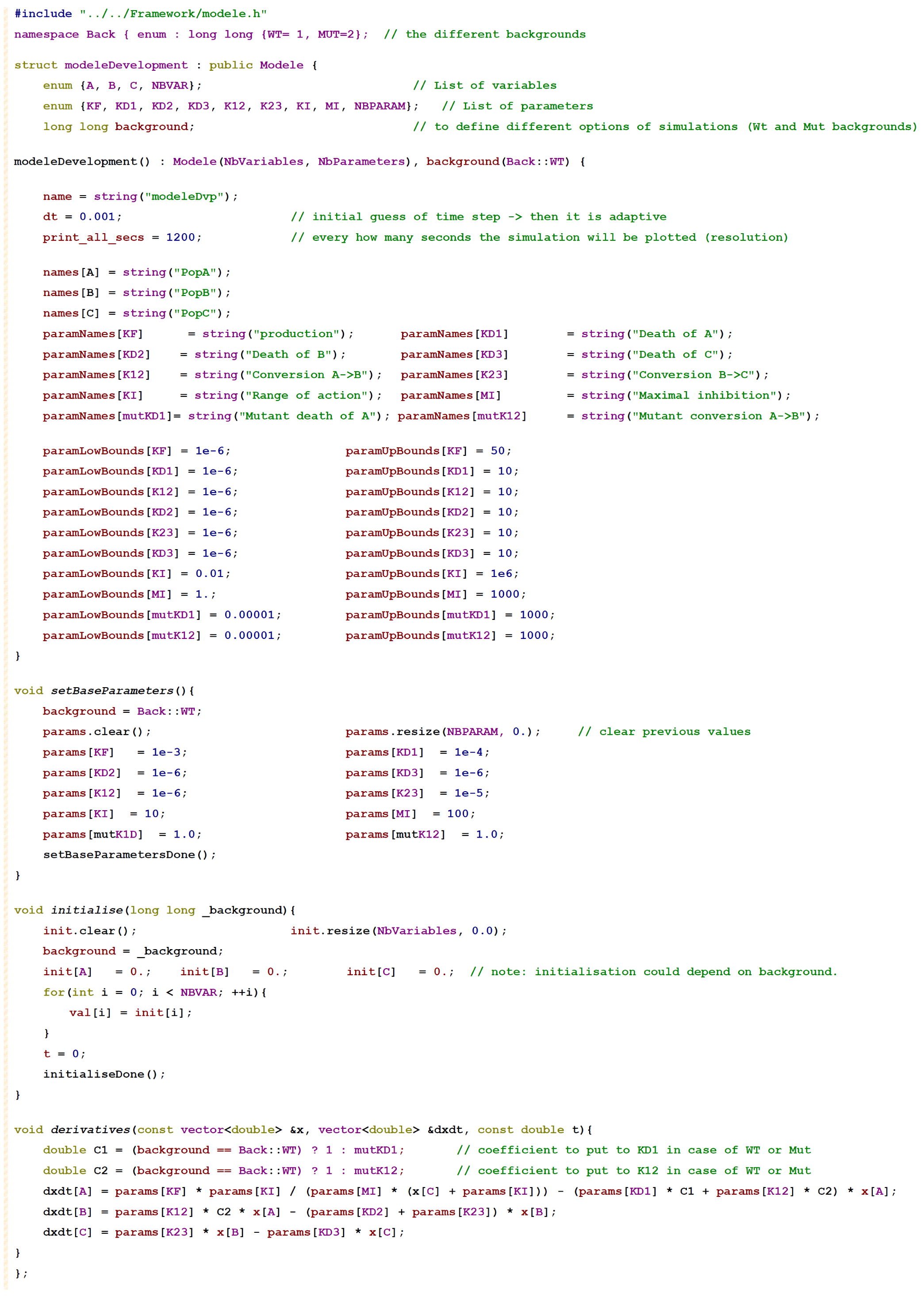
Code for the modele of 3-stages cellular development with feedback.

**Figure 6:**
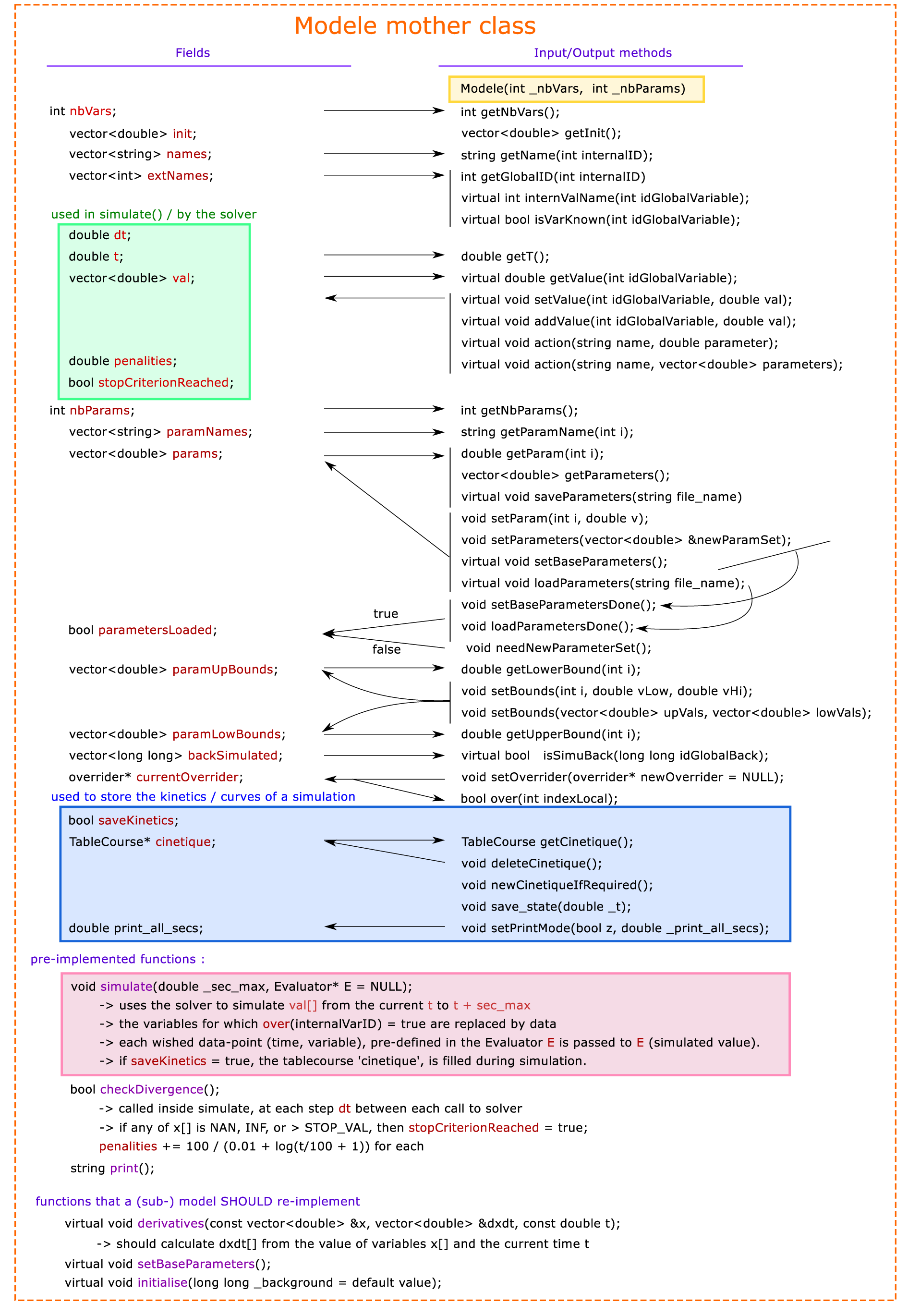
Memory fields provided by the Modele mother class, and the input/output functions to access them. Most fields contain information filled by the sub-model at instantiation (names, extNames, paramNames, paramUp/LowBounds, etc.) The fields inti and params contain information to make one simulation. The fields dt, t, val, penalties and stopCriterionReached are used by the solver to perform a simulation. currentOverrider contains the information, for each variable, whether it should be overriden by a predefined curve, and if yes, it contains the function of that curve. Finally, cinetique is used to store the kinetic data for each variable during a simulation, every print_all_secs seconds, if the flag saveKinetics is true.

A submodel should only implement four functions:

1. a constructor that fills the fields names, extNames, paramNames, paramUpBounds and paramLowBounds, and calls the Modele mother constructor by providing the number of parameters and variables. As the variables and parameter values are manipulated as vector<double>, it is suggested to define two enum to name the position of variables or parameters inside the vectors by a real name.
2. a function derivatives according to the virtual function of the mother class. If there are NV variables, the vector x and dx are of dimension NV (each dimension is a variable). The vector x is the current value of each variables, dxdt is the output vector to be filled according to the ODEs, and t is the current time, if needed. It is suggested to name the variables using an enum, in which case the value of a variable will be *x* [*name*_*v*_ *ariable*_*i*_ *n*_*e*_*num*].
3. a function setBaseParameters that puts a default set of parameter values into params. It is recommended to call the function ‘setBaseParametersDone()’ of the mother Modele class, to confirm that a set of parameter values has been set.
4. a function initialize that puts the intial value of each variable (inside init), depending on the current parameters. Additionally, it can restart the time of simulation t and apply the initial values to the variables. It is possible to give an option to initialize, here called background, such that the same model could be run or initialized under different conditions while using the same set of ODEs. It is recommended to call the function ‘initialiseDone()’ of the mother Modele class to confirm that initialization has been done.

The model presented as example in section 1 can be written as the following C++ subclass (Figure 5). The example ‘Development.cpp’ shows a more complex version of this model (see the folder Examples).

Additionally, many functions are already provided inside the Modele mother class (see figure 6). First, the virtual functions re-implemented by the sub-model can be called to create an instance of the sub-model and to initialize it. Then, the simulate function solves the ODEs of the sub-model for _sec_max seconds. If an evaluator E is provided (containing time-points and variables to record), see below, the simulation will automatically make breaks at each desired time-point and fill the evaluator E with the simulated data at these time-points. The overrider class stored in the model (currentOverrider) contains information whether some variables should be replaced by kinetic interpolated data. The function simulate override the appropriate variables at each time-step before calling the solver and to erase them afterwards before the solver computes the simulation error. InitializeDone has to be called by initialize from the submodel in order to automatically clear kinetics. Further, the function checkdivergence is called by the simulate() function to stop a divergent simulation as early as possible, and can be modified if wanted.

In order to run it from the interface, two more steps will be required: defining different experimental set-ups and loading experimental data. From a C++ script, basic functions can also be run directly from the model, for instance:

~~~
Modele* M = new modeleDevelopment ();
M->print();
M->setBaseParameters();
M->initialise();
M->simulate(36000); // simulates for 36000 seconds. By default, nothing stored
M->setPrintMode(true, 1200) // Now, will save every 1200 seconds
M->initialise();
M->simulate(36000); // simulates for 36000 seconds
TableCourse TC = M->getCinetique()
cout << TC.print() << endl;
~~~

### 3.3 Loading experimental data and getting the cost of one simulation

A first way to load experimental data is to use text files, where data are separated by tabulations. An example of data file and how to load them with the class TableCourse is shown in Figure 7.

**Figure 7:**
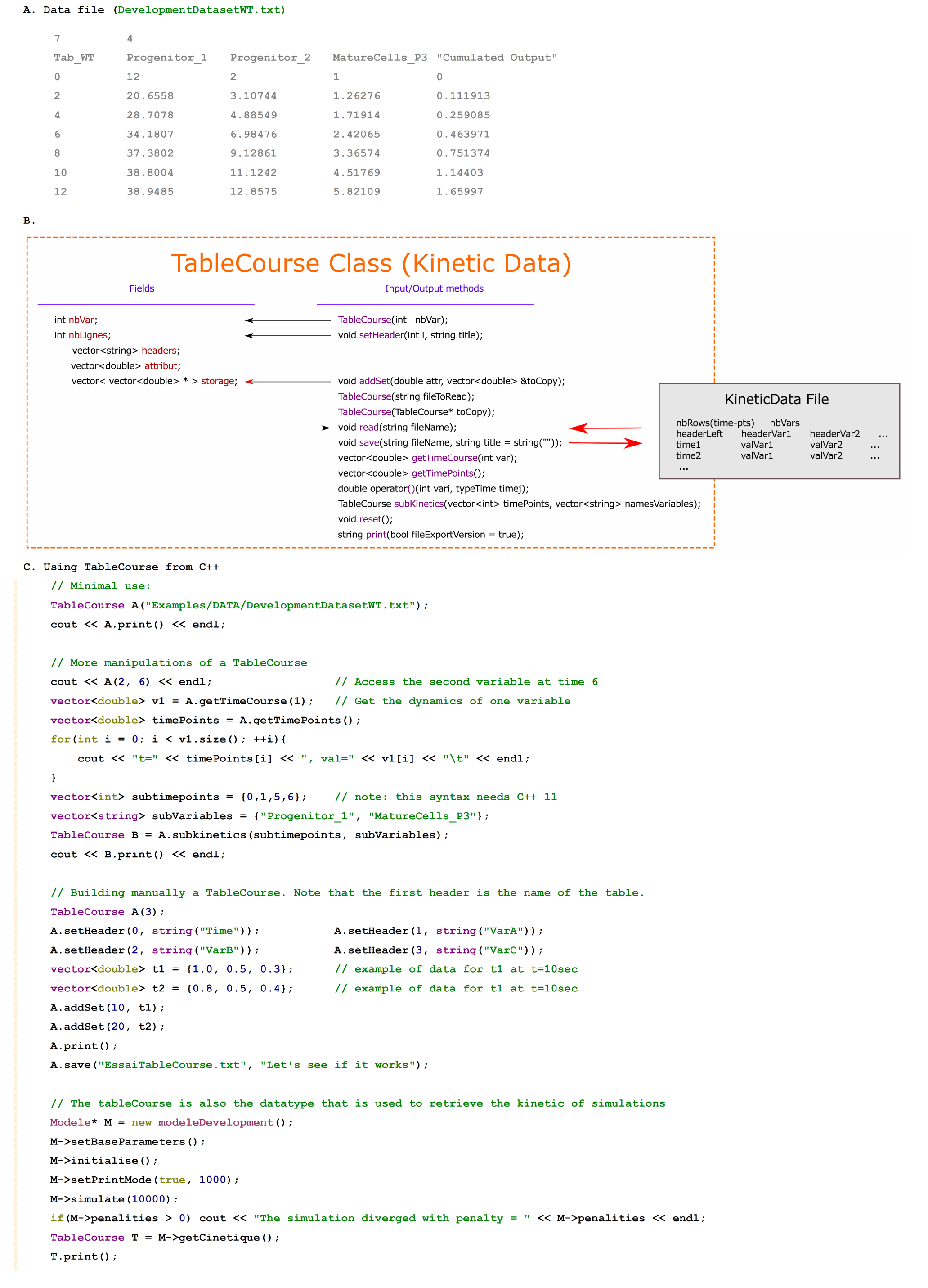
Loading experimental data from text files. A. File format. The file should specify: the number of datapoints NT and the number of variables NV, then give the name of the table and of all variables, without space or by using “ “, and then, for each time-point, the time and the value of each variable. Note: use ‘.’ for decimals and not ‘,’. In this example, 7 time-points and 4 variables, and the name of this table is ‘Tab_WT’. B. Structure of the TableCourse class, provided to read those files. C. Examples of use.

Optimization requires to perform a huge number of simulations, and it would be time consuming to store detailed kinetics for each simulation and each variable. Therefore, the list of experimental datapoints can be defined before simulations/optimizations. Inside the function simulate(), the solver will automatically stop at these time-points and only the the simulated values at this time-points will be stored, this is done by using the ‘Evaluator’ class. An ‘Evaluator’ stores a list of ‘datapoint variable experimental mean experimental standard deviation (optional)’, and can also store simulated data for these datapoints in order to get a ‘cost’ comparing the simulation and the data.

The main functions to load data files into an evaluator and to retrieve the cost of a simulation are shown in Figure 8. Experimental data can be given to Evaluators either from a textfile (see Figure 7) or manually using getVal() ^1^. In both cases, it is advised to give the mapping between the name of the variable in the text files and a ‘global ID’ or enum of this variable ^2^. Finally, once an Evaluator is filled with experimental data, the function recordingCompleted() has to be called.

**Figure 8:**
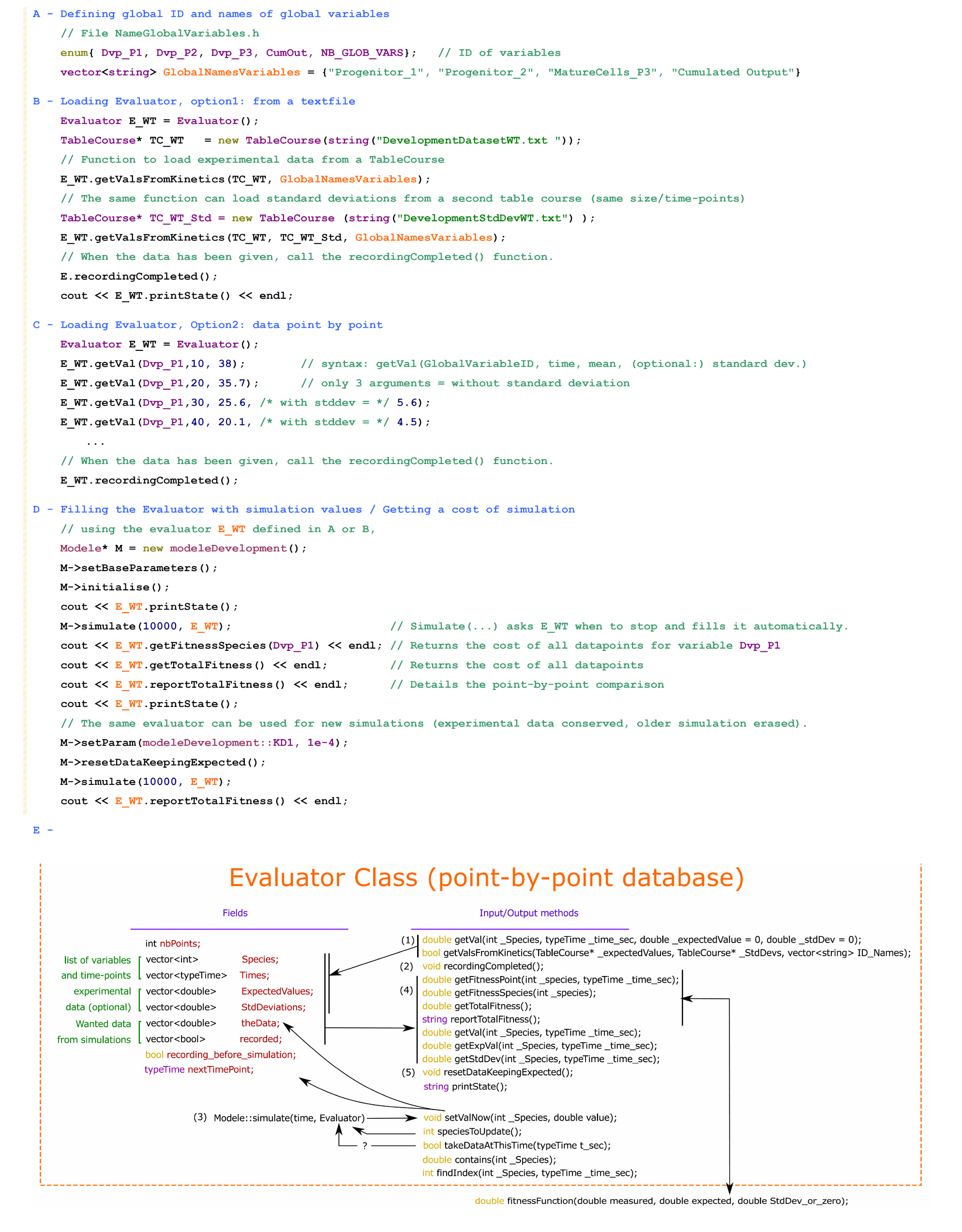
Use of the Evaluator class. A. For using an Evaluator, it is required to link the name of variables to an ID (integer). B. Loading experimental data in Evaluators from text files, by using the list of variable names (see A). C. Manual loading of an evaluator. D. In order to get the cost of a simulation, an evaluator has to be loaded first, then passed to the function simulate that will fill the Evaluator, and then the cost or fitness functions of the evaluator give the combined cost of each time-point and/or variable. E. Fields and methods of the Evaluator class. The numbers represent the order of events during a simulation and cost calculation. After giving datapoints (1) and setting recordingcompleted (2), the simulate function will get informations on which next time point to stop and and which variable value is requested. Then, when this time-point is reached, the simulate function gives the simulated value to the evaluator via setValNow (3). After the simulation, the fitness functions can be used to evaluate the cost of the simulation (4) by calling the fitness function (customizable). Finally, the evaluator has to be reset (5) before the next use. Many simulations can be performed successively by repeating the steps 3-4-5.

During a simulation, an Evaluator is filled with simulation values via the function setValNow(variable, time). This is performed automatically inside the Modele::simulate() function. However, it is possible to do it manually by following the steps: 1/ when is the next stopping time-point where simulation data is required ? by calling takeDataAtThisTime(t). The first value *t* equal or exceding the next ‘stop-ping’ time-point will raise *t rue*. 2/ At the current time-point, what is the next variable requested by the evaluator ? speciesToUpdate() returns the ID of the wanted variable. 3/ Now, setValNow(variable, value) can be called to provide the simulation value of this variable at the current time-points.

### 3.4 Defining the experimental settings

Additionally to the ODE model, a set of experimental conditions has to be defined. This is done as creating sub-classes of the Experiments subclass. An experiment takes a Model sub-class and defines a number NC of different conditions. Each condition states what has to be simulated: from which initial conditions, how long, and which perturbations of the system, in a way to reproduce the experimental data or to simulate predictions. Therefore, the same experiment / conditions can be performed indifferently with different models. Additionally, each experiment allow to store experimental data in its ‘Evaluators’, one evaluator per condition (stored in the field VTG, ‘Values To Get’). The main fields of the mother class Experiments and the design that a sub-class should follow, are shown in Figure 9. An experiment (subclass of Experiments mother class) has to:

**Figure 9:**
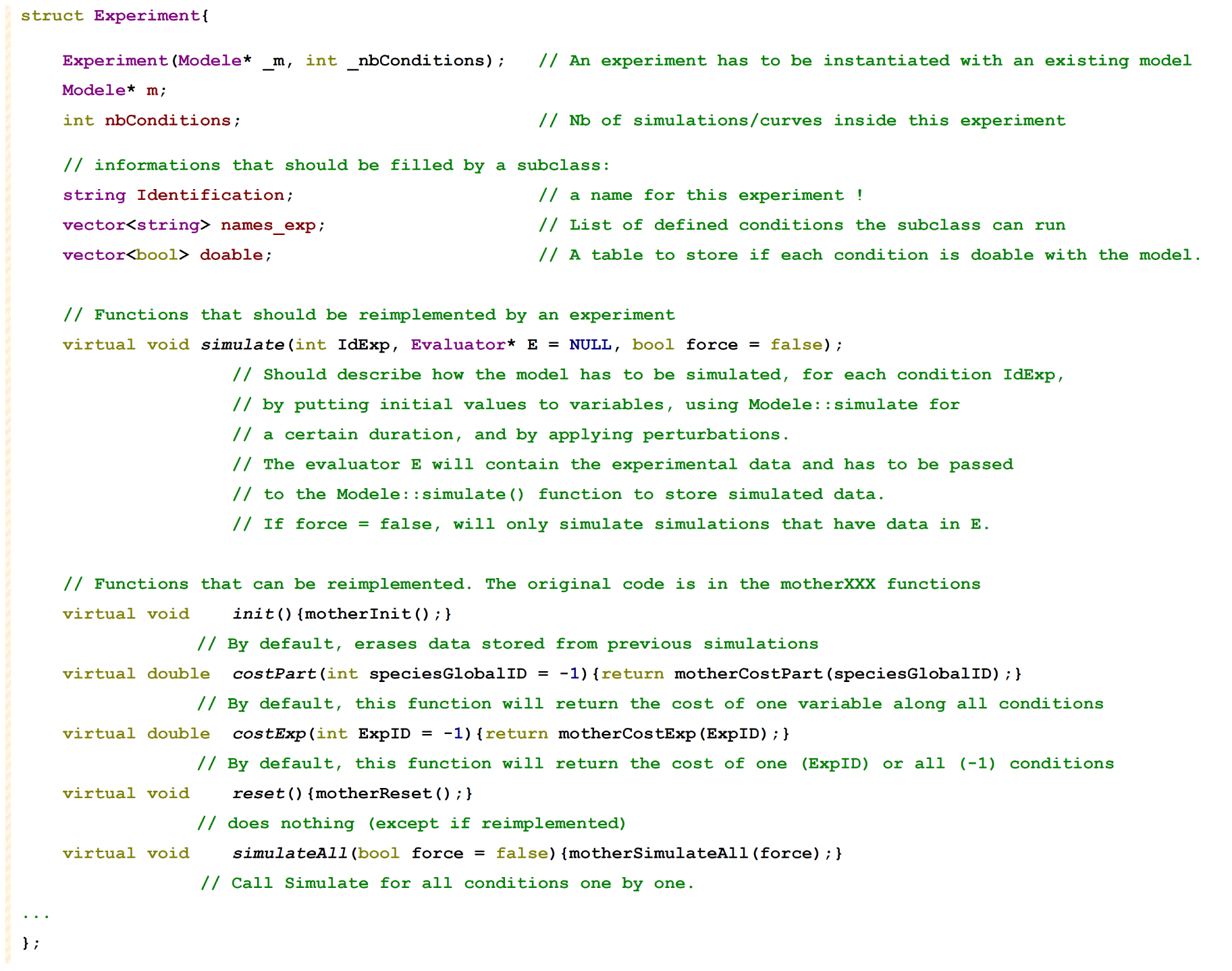
Main structure of the experiment mother-class, defining how an experiment should be structured. An experiment only needs to define a number of conditions, their name, and to implement a simulate function able to simulate each condition from the model. Most functions for handling data and performing grouped analyses on all conditions (cost functions, etc) are already implemented but can be customized by rewriting the virtual functions in the subclass.

1. implement a constructor that calls the Experiments constructor, giving the model m (that has to be already instanciated first). Inside the constructor, give the name of all the experiments in names_exp and, if some models do not fit some experiments) update the table ‘doable’, which says for each experiment if it can be simulated. For instance, by asking the it if he can simulate variables or backgrounds.
2. implement a function that does simulate each different conditions
3. additional functions (init, getCost …) can be reimplemented

A minimal code to represent the experimental setting of the example of section 1 is shown in Figure 10, and the details of the available functions is depicted in Figure 11.

**Figure 10:**
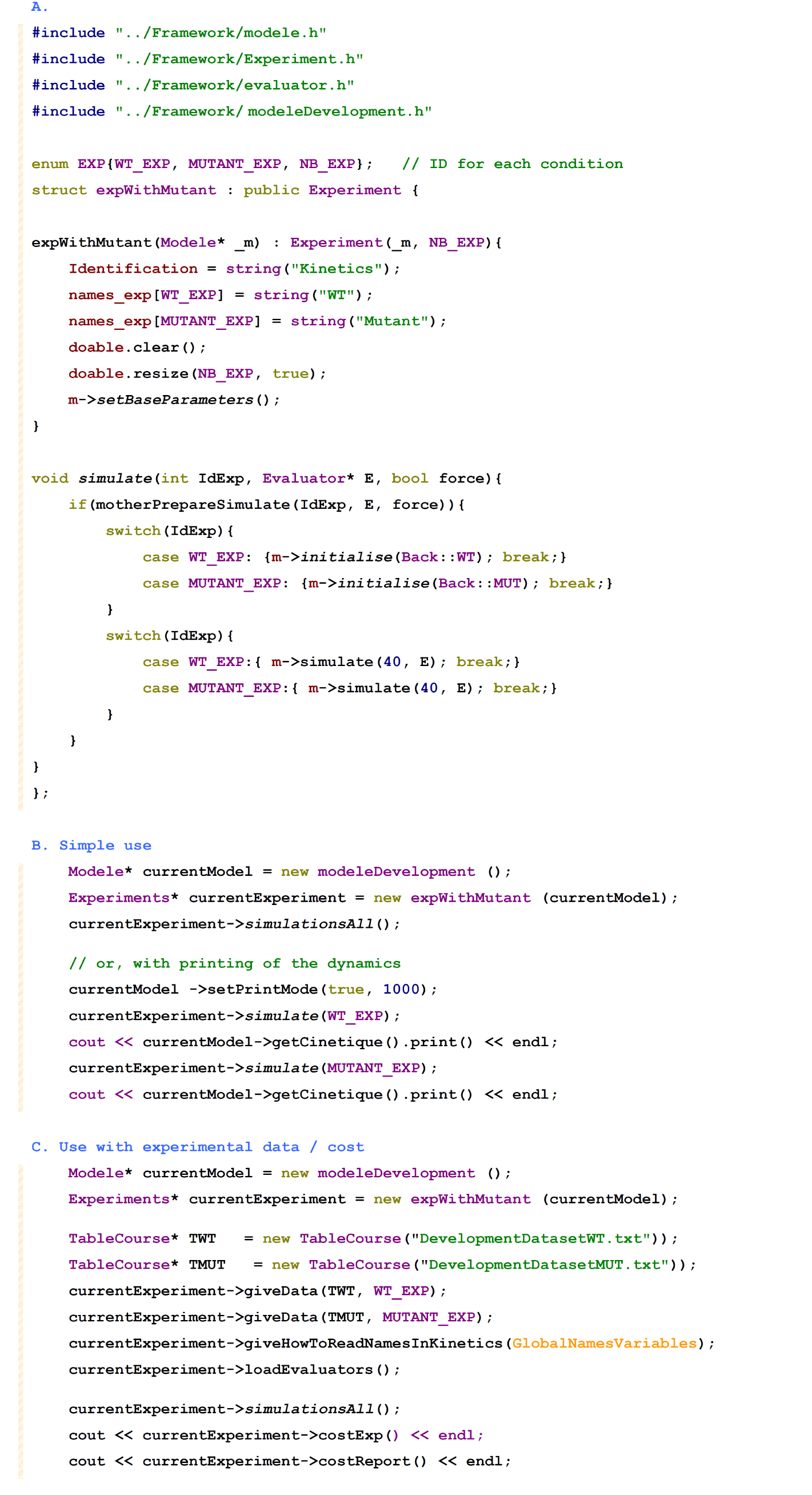
Defining and using experiments. A. Example of simple code to represent the experimental conditions of the example shown in Figure 3. B. Example of syntax to instantiate a new experiment from a model and run a simulation. C. Example on how to load experimental data to an experiment directly, to perform a simulation and to retrieve a cost. The experiments has pre-defined evaluators (VTG[]), one per condition, and the function ‘giveData’ allows to automatize the commands shown in Figure 8 to load the evaluators, from the ID of each condition.

**Figure 11:**
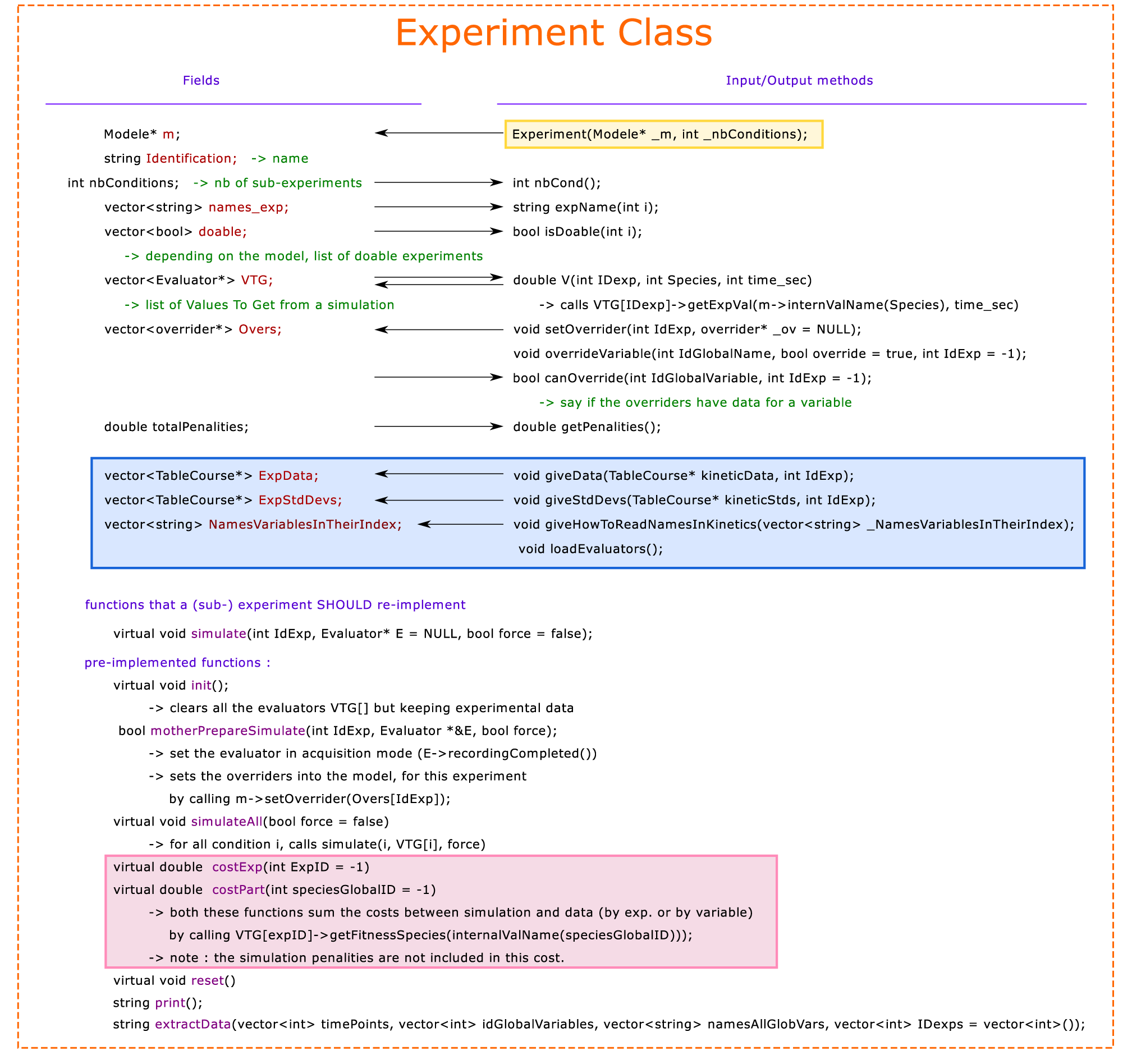
Details of the fields and functions provided by the experiments mother class. Depending on the model, some conditions might not be doable, and can be notified inside doable[ID_condition] = true or false. The vector VTG stores an evaluator for each condition. The preimplemented functions init() and motherPrepareSimulate() automatically prepare the evaluators for a simulation. The experimental data can be stored in ExpData and ExpStdDevs using the function giveData() and giveStdDevs(). Finally, the field Overs stores a list of curves to be used to replace certain variables [see section 4]. After a simulation is performed, the function costExp() and costPart() request the Evaluator to give a cost for each condition.

### 3.5 Running simulations or optimizations from the graphical interface

The interface only requires an existing experiment to be instantiated before, and will be launched by calling a new simuWin class from the experiment. The commands to start the GUI from a model / experiments are shown in Figure 12. Alternately, it is possible to perform optimizations in a script (without interface) by calling a new manageSims instead. The functions to optimize a list of combinations (of which parameters to optimize separately), and to merge the parameters from each fittings are included in Figure 12, as an example. The list of functions provided by the manageSims class are shown in Figure 13.

**Figure 12:**
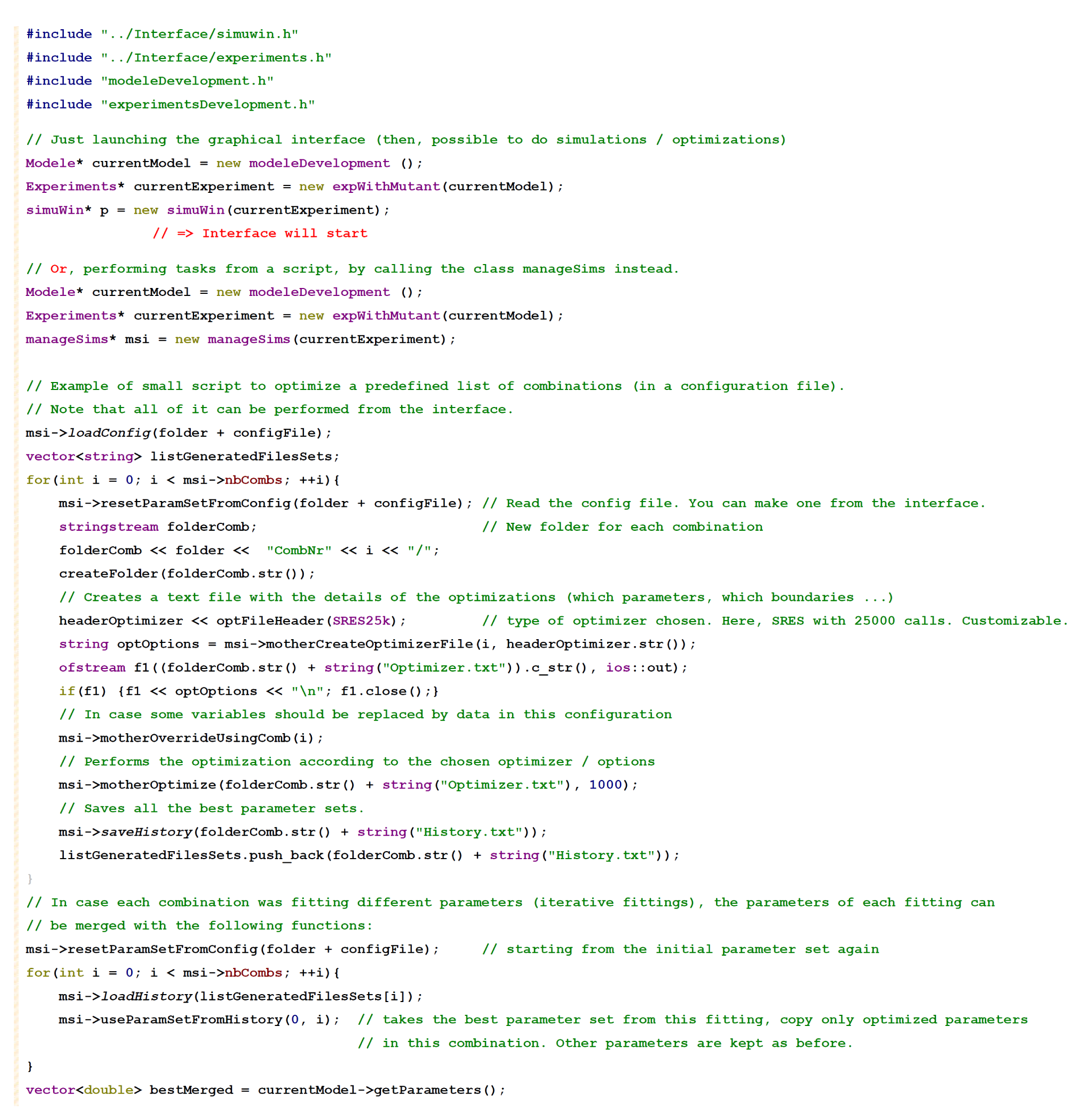
Performing simulations / optimizations. Syntax to call a graphical interface from an experiment, or to perform iterative fittings of different combinations of parameters, as defined in a configuration file. Configuration files are text-files that can be created from the interface.

**Figure 13:**
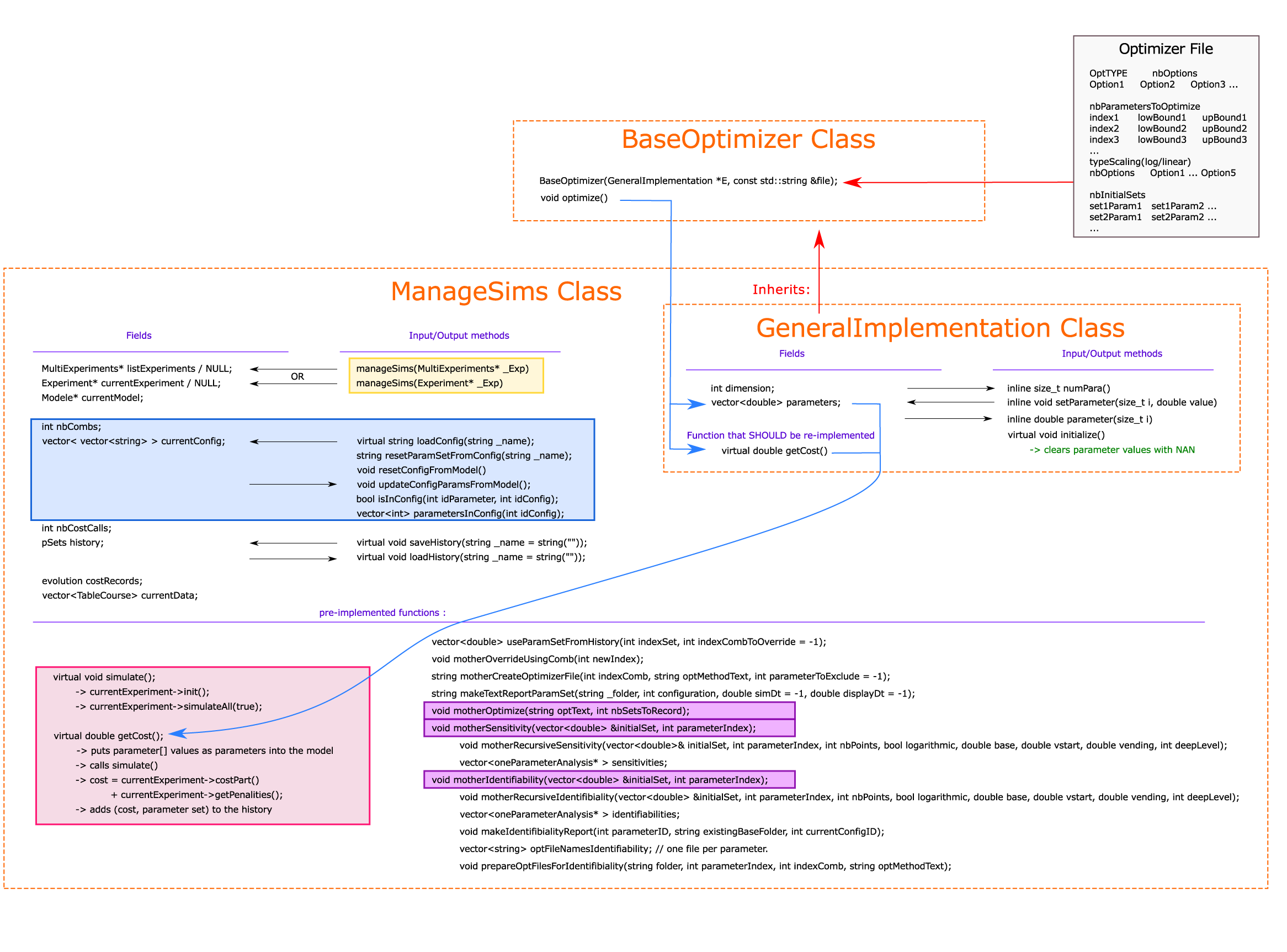
Design and functions in the manageSims function, and interplay with the optimization classes. It is constructed based on an existing experiment (containing an existing model). A configuration (list of combinations of which parameters to optimize) is stored in currentConfig, and can be loaded or saved from text files. History saves the list of best parameter sets from the last optimization. The class manageSims itself is defined as a sub-class of a ‘GeneralImplementation’, i.e. a class with a list of parameters and a getCost() function. Therefore, any optimization toolbox can be used to perform fittings, by taking the created manageSims as a ‘generalImplementation’. The BaseOptimizer class provides already implemented optimizers, and parse the informations written into the optimizer options files. Finally, the graphical interface inherits from the manageSims class, and is just a wrap-up that calls the functions of manageSims according to the choices of the user.

**Fitting results for the example model.** Figure 14 shows typical results of fitting using MoonFit. On the left side, the model without feedback is unable to recapitulate a decrease in T1 or T2 over time, and the curves with minimal cost lie in between the datapoints. However, on the right side, fitting the model with feedback allows to fit the data with a fair quality, even with only two parameters changing between the WT and mutant background: *K*_*D*1_ and *K*_1 − 2_. In particular, it raises an increase death of the T1 population and a lower transition. It doesn’t prove that the model is the accurate description of B cell development. Indeed, the model is likely to be overfitted or non-identifiable (too many parameters), and more experimental data or a validating dataset could help confirm the validity of the model. However, it shows that the proposed interpretation of the authors is indeed compatible in a quantitative way with the phenotype of the deficient background, which is already an important step.

**Figure 14:**
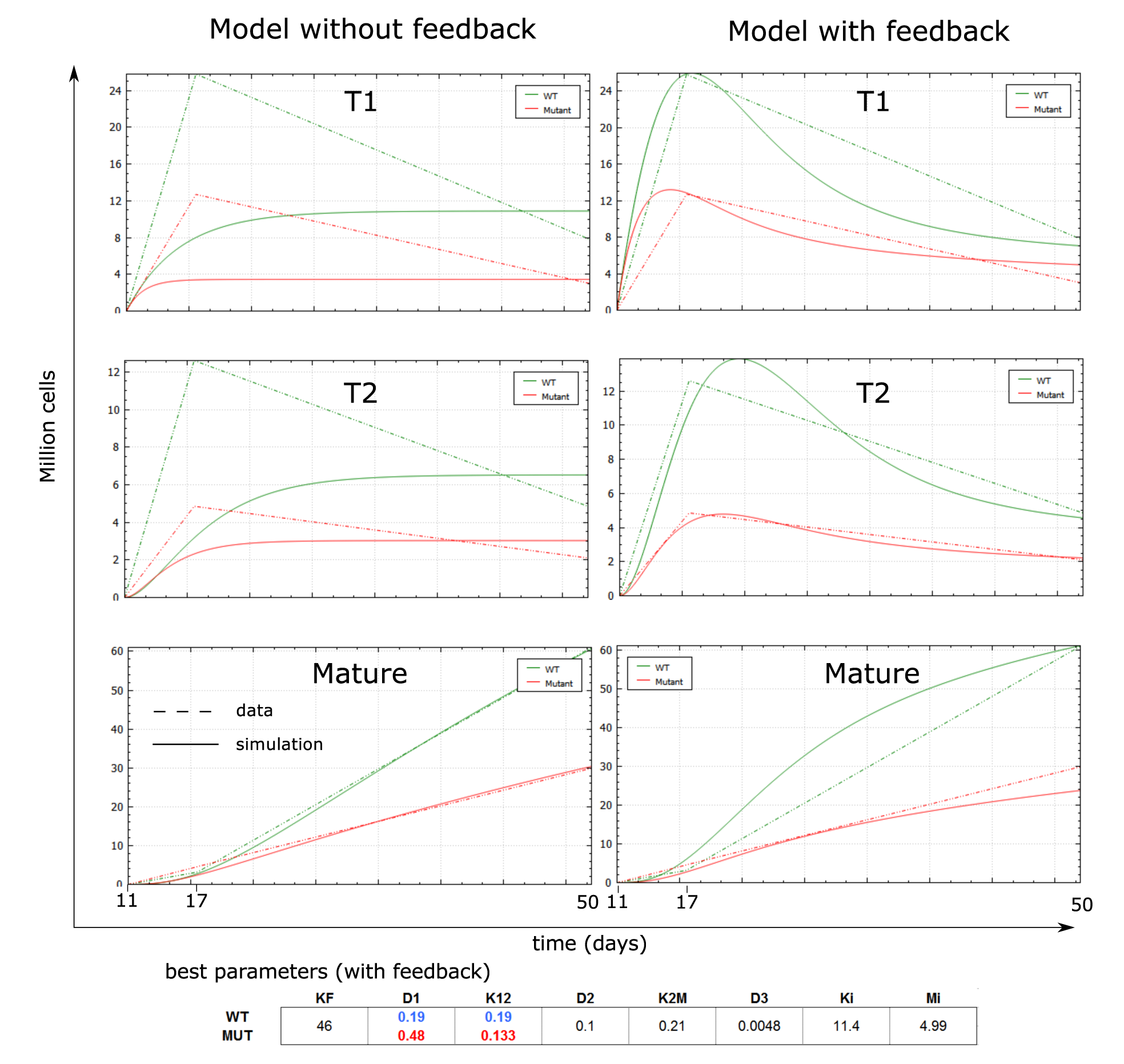
Fittings of the B cell development example/dataset with the two proposed models. The dynamics of best fitted simulations for the model without feedback (left) or with feedback (right) are depicted for the three populations of interest. Dashed curve represent the datapoints shown in Figure 3, and solid, smooth lines represent simulations.

## 4 Advanced options

### 4.1 Performing iterative fittings / override variables from data

Iterative fitting is a strategy that consists into simulating only a few variables while replacing other variables by their experimental dynamics during the simulations [20].

For instance, in the context of Gene Regulatory Networks simulating both mRNA and protein levels,if all the protein levels are replaced by an interpolation of the measured dynamics, then the mRNA variables become independent to each-other (they only depend on protein levels for their regulations usually), and can be fitted separately. This technique can be important when the data shows latencies between mRNA and proteins for instance, and can be used to quantify them. Further, iterative fitting is advantageous in term of speed and quality of fitting [20].

Using the interface, iterative fitting is performed by creating combinations (one column per combination), containing which parameters to optimize (top part) and which variables to replace by data (bottom part). A complete iterative fitting would consist into creating disjoint combinations, where each combination fits a subset of parameters, and then a first global parameter set would be obtained by merging the values from each separate optimization. Then, a global optimization 20% to 50% around this ‘merged’ parameter set would be advised, without any overriding. Note that, for an experiment, a variable can be overrided only if a it has been given data for all the conditions. Figure 16 details how to provide experimental data for overriding, and the code presented in Figure 13 is suitable for doing iterative fittings and merging the parameter sets, from a given configuration.

### 4.2 Changing the cost function

The cost function is defined inside Evaluator.cpp. The common square and logarithmic cost functions can be used by having one of the following options uncommented in Evaluator.h: *SQUARE*_*COST, SQUARE*_*COST* _*ST D, LOG*_*COST* or *LOG*_*COST* _*ST D*. Generally speaking, the function fitnessFunction() is called for each datapoint and can be modified or replaced by another kind of cost.

For mode complicated cost-functions that would add coefficients to different experiments for instance, we would suggest to re-implement the getCost() function of the experiment to modulate the weight of the cost of each experiment. Also, a class ‘MultiExperiment’ is defined inside Experiment.h/cpp that allows to pool multiple experiments inside one fitting, with a different cost each.

### 4.3 Using a combination of backgrounds

The ODE models are only defined with one option (background) to be initialized differently. If a model can simulate several backgrounds, one might want to simulate according to a combination of backgrounds, such as a double deficient background. An example is given in Figure 15.

**Figure 15:**
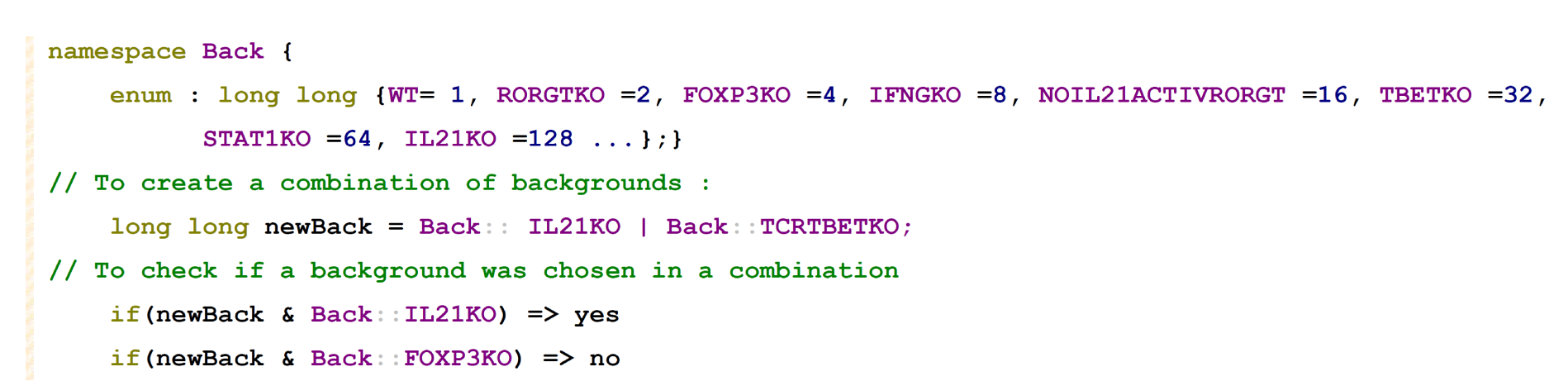
Using combinations of backgrounds.

**Figure 16:**
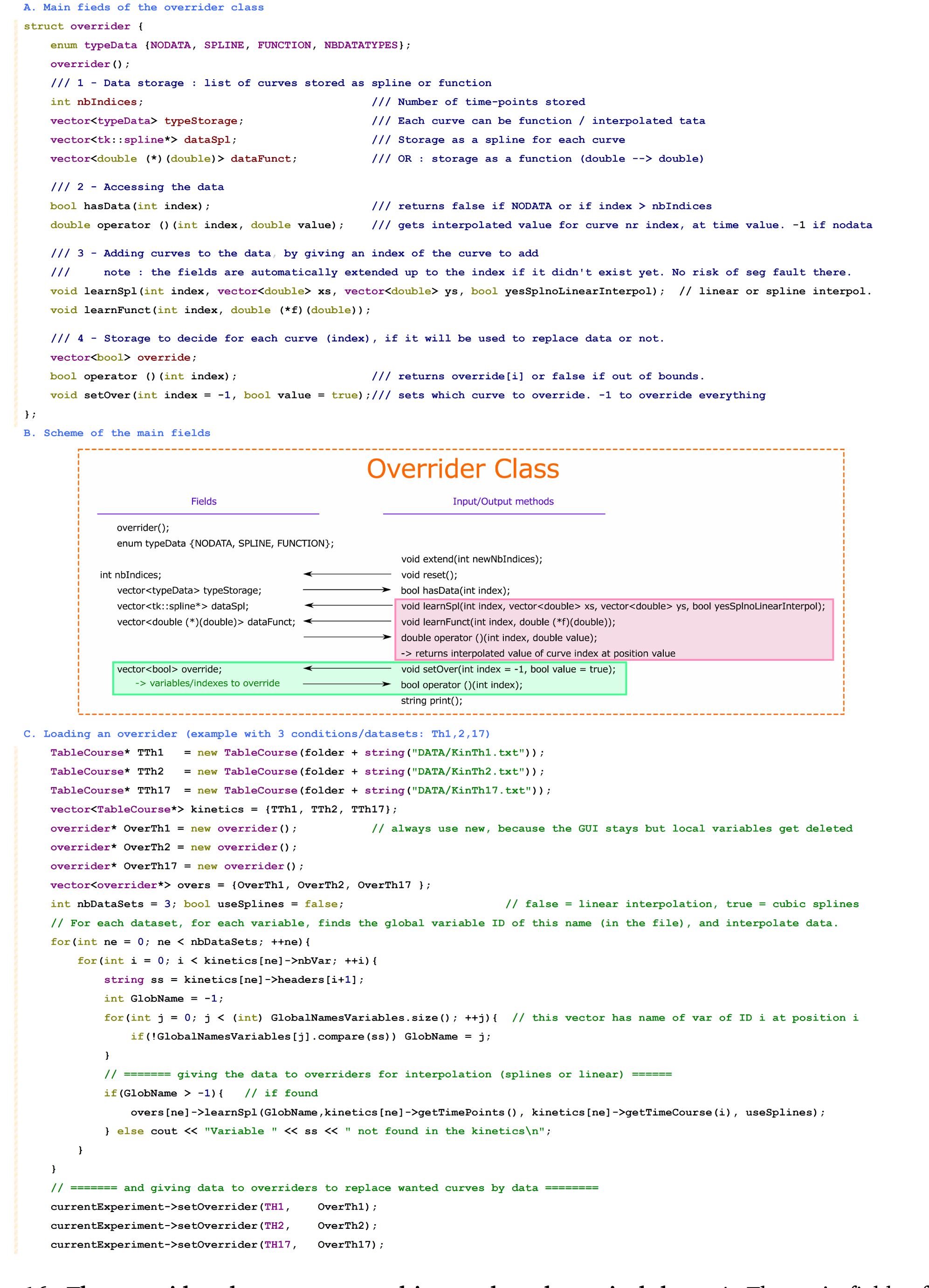
The overrider class to store and interpolate dynamical data. A. The main fields of an overrider. It can store a list of either interpolated curves (see the tk::spline library), or a function from double (time) to double (value). B. Scheme of the main functions. C. Example of code to load an overrider, by interpolating the curves of each variables one by one and giving it to an experiment.

### 4.4 Changing the solver

The function Modele::simulate calls the built-in solver with the common syntax used by solver libraries like boost. A scheme of the simulate function and how the overriding of variables by data is performed is shown in the supplements in Figure 17. It includes the example how to use the boost solver instead of the original one.

### 4.5 Changing the optimization method

The list of provided optimizers is shown in the supplements in Figure 18. There are two ways to choose an optimizer.

**Tuning a provided optimizer.** When using one of the provided solvers, the interface simuWin (or manageSims), will always create an ‘optimizer file’ in the current folder, before starting an optimization. This text file contains the type of optimizer requested (ex: SRES, GeneticGeneral), the list of parameter indexes and boundaries, and the choice to optimize in logarithmic or linear scale. During this process, a ‘header’ of the optimization file for the optimizer options is always taken as argument, and can be defined manually.

**Using an external library.** When a fitting problem is started as a simuWin or manageSims, these both classes are designed as sub-classes of a ‘baseOptimization problem’, meaning a simple struct with a list of parameters and a cost function. So any external class can use simuWin or manageSims as a blackbox. Even better, by giving a simuWin to an optimizer, the parameter choices of the optimizer can be monitored in real time together with the evolution of cost. In order to give the options to the external optimizer, it would be required to read the optimizer text file generated before starting the optimization. It would be advised to use the class ‘baseOptimizer’ to read this file and then to give the options to the external optimizer. Alternatively, by making the external optimizer inherit from baseOptimizer, it can read those files by itself.

### 4.6 Automated generation of models

The file generate.cpp contains both a graph class to add different kinds of interactions between different factors as a ‘matrix’. Then, a running code is creating the C++ code for the model class defined from that graph, using hill functions. Further, a function subgraph allow to automaticall generate all possible subgraphes of a network, with a wanted number of interactions, and each sub-network can then be generated as a different model.

## 5 Discussion

We presented here a software that brings together home-made C++ defined ODEs with a user-friendly graphical interface, a solver and a few optimizers.

It is particularly interesting in the case of models that need to be refined or programmed under specific needs that do not necessarily match the standard allowed ODE definitions from big tools or library. Also, it allows to compare different and home-made optimizers under the same interface, and it allows to perform iterative fittings [20], which is not yet widely used nor available.

When a network is not suited to a dataset, parameter optimization consistently fails, and a lot of manual fitting and refinements needs to be performed. The fact to have an interface to directly resimulate new parameter sets and visualize the curves can help to save time.

The modular structure of the code was designed to be re-used independently if needed. For instance, the Modele.cpp class can be used alone to solve ODE. The interface could be used with a different kind of experiment as well, and so on. One could also modify the modele class to simulate other types of models such as agent-based, that do not need a solver but still follow a description of parameters and experiments, in which case the simulate() function can be reimplemented by another kind of time simulation.

One limitation however, is that the optimizer is designed to be completely independent and to consider the model/experiments as a blackbox with a number of parameters and their boundaries. It means that complex optimizers that use additional information from the ODEs like gradient methods based on the Hessian matrix can not easily be run in this context. Also, experimental data sometimes include scaling or normalizing factors, which is not provided here, and would need to be performed manually when comparing a simulation to a dataset.

## Footnotes

1 The function Evaluator getVal() can be used both for loading experimental data (before calling recordingCompleted()), and for retrieving the simulated value for that point (afterwards). In case of a customized cost function based on getVal(), a first call of this function loads the evaluator with the desired time-points, while, after recordingCompleted, the very same function can be used to compute a cost.

2 In case it is wished to develop models with different complexity for the same biological process, the internal variables of each model can be linked to the global ID of that variable (field Modele::extNames), and in general, any action performed on models is done using the global ID of variables (for instance using setValue).

